# In silico analysis of imprinted gene expression in the mouse skin

**DOI:** 10.1101/2020.06.04.134031

**Authors:** Alexandra K. Marr, Sabri Boughorbel, Mohammed Toufiq, Mohammed El Anbari, Aouatef I. Chouchane, Tomoshige Kino

## Abstract

Imprinted genes help mediate embryonic cell proliferation and differentiation, but their roles after birth are far less well understood. A subset of 16 imprinted gene network (IGN) genes is expressed at higher levels in stem cell progenitor cells of adult skeletal muscle and epidermis compared to their differentiated counterparts. While these genes function in muscle regeneration, their role in the skin is poorly understood. We assessed the expression profiles of these 16 IGN genes in publicly available datasets and revealed elevated expression of IGN genes in the telogen and early anagen phases in mouse skin. We also identified IGN genes among a list of previously identified hair cycle-associated genes. Furthermore, our results suggest that IGN genes form part of a larger network and function predominantly as upstream regulators of hair cycle-regulated genes. Based on these *in silico* data, we propose a potential novel role of these 16 IGN genes as upstream regulators of hair cycle-associated genes. We speculate that IGN gene dysregulation participates in syndromes characterized by an impaired hair cycle. Thus, IGN gene expression might serve as a point of therapeutic intervention for patients suffering from cutaneous pathologies such as common hair-loss disorders.

## 2. Introduction

Genomic imprinting is an epigenetic regulatory mechanism that confers expression of selected genes from one parental allele, and thus, is independent of classical Mendelian inheritance ^1^. Genomic imprinting is established in parental germline cells, and is maintained throughout mitotic cell division in somatic cells ^2^. Thus far, 85 and 95 murine imprinted genes have been reported in the COXPRESdb and Gemma databases, respectively ^3^. In mammals, imprinted genes are commonly identified in clusters located on specific chromosome regions ^4^. The process of gene imprinting involves coordinated DNA and histone methylation, whereas the mechanisms underlying the selective targeting of a particular set of genes are largely unknown ^5^. Altered expression of imprinted genes has been associated with the development of various pathological conditions in humans, including obesity, diabetes mellitus, muscular dystrophy, mental disability and neoplasms ^1^.

Imprinted genes are functionally distinct, but most are involved in controlling the transition of cells between their quiescent, proliferative and/or differentiated states during fibroblast cell cycle withdrawal, adipogenesis *in vitro*, and muscle regeneration *in vivo* ^1^. These genes function cooperatively in the regulation of specific biological pathways by forming co-expressed networks ^1^. One of these subnetworks comprises 16 imprinted genes (hereafter referred to as the IGN genes) [cyclin-dependent kinase inhibitor 1C (*Cdkn1c*), decorin *(Dcn)*, delta-like non-canonical notch ligand 1 (*Dlk1*), glycine amidinotransferase (*Gatm*), *GNAS* complex locus (*Gnas*), growth factor receptor-bound protein 10 (*Grb10*), imprinted maternally expressed transcript (*H19*), insulin-like growth factor 2 (*Igf2*), insulin-like growth factor 2 receptor (*Igf2r*), maternally expressed gene 3 (*Meg3*), mesoderm-specific transcript (*Mest*), necdin (*Ndn*), paternally expressed gene 3 (*Peg3*), PLAGL1-like zinc finger 1 (*Plagl1/Zac1*), sarcoglycan, epsilon (*Sgce*), and solute carrier family 38 member 4 (*Slc38a4*)]. A hallmark of gene networks is their ability to alter the expression of member genes in response to environmental changes. Several imprinted genes modify the expression of others within the same gene network. For example, *Plagl1/Zac1* controls embryonic growth by influencing the expression of *Igf2, H19, Cdkn1c* and *Dlk1* ^5^. In addition, Gabory *et al*. demonstrated that *H19* gene knockout alters the expression of *Igf2, Cdkn1c, Gnas, Dlk1* and *Igf2r* in mice ^6^. The majority of IGN genes are expressed at high levels during embryonic and early postnatal life, but are silenced in the adult, except in muscle satellite cells, hematopoietic stem cells and skin stem cells ^5,7^. In mouse skin tissue, it has been reported that *Cdkn1c, Dlk1, Grb10, H19, Igf2, Mest, Ndn, Peg3* and *Plagl1* are expressed at higher levels in epidermal stem cells compared to those in non-stem cells (keratinocytes) ^7^. While imprinted genes have been shown to play a role in muscle regeneration and hematopoiesis, their functions in skin tissue are poorly understood ^8,9^; thus, this was the focus of our study.

Skin stem cells are multipotent adult stem cells that can self-renew and differentiate into multiple cell lineages to form the different layers of the skin as well as the hair follicle. The cyclic activity of hair follicles organizes the growth and renewal of hair. During its life span, hair undergoes growth, degeneration and regeneration in concert with the activation and quiescence of epidermal stem cells located in the bulge of the hair follicle ^10,11^. The cyclic activity of hair growth is divided into the anagen (growth), catagen (regression), and telogen (resting) phases ^12^. Follicular stem cells are maintained in a quiescent state during the telogen phase. Once activating signals are received from upstream regulatory systems, a new cycle of hair growth is initiated (anagen phase) ^11,13,14^. After the active growth phase, proliferating matrix cells in the hair follicles are induced to undergo coordinated apoptosis (catagen phase) ^12^. Following the catagen phase, the hair follicles eventually undergo transition to the telogen phase, during which hairs are no longer produced due to inactivation of the follicular stem cells ^12^.

In this study, we hypothesized that IGN genes play important roles in skin/hair biology throughout life (after birth) in addition to their well-known activity during embryonic/fetal growth. Thus, we aimed to identify the potential function of the 16 IGN genes in mouse skin tissue after birth by reanalyzing publicly available transcription profiles. Here, we exploited the curated large-scale datasets held in the NCBI GEO Profiles database. This public repository contains more than 70,000 transcriptome data series, with over 1.8 million individual profiles ^15^ and offers the option to examine the abundance of individual genes determined in hundreds of ‘omics’ studies. To identify datasets with changes in the abundance of IGN genes, we initially selected *H19* as a representative of imprinted genes because it is known to influence the expression of several other genes in the IGN ^16^. We then assessed the expression profiles of all 16 network-forming imprinted genes in datasets of interest. Using this strategy, we aimed to identify potential gaps in knowledge about IGN genes in skin biology based on changes in the corresponding RNA abundance with the long-term goal of supporting the development of novel therapies for skin disorders and hair-loss conditions.

## 3. Results

### 3.1 Identification of differential expression of *H19* in different stages of the hair cycle in mouse skin

To study the expression of IGN genes in mouse skin, we searched the NCBI GEO databank for a dataset that includes untreated, unaffected mouse skin samples. Using the search term ‘skin AND C3H/HeJ’, we identified dataset GSE45513 which contains three samples of skin transcription profiles from 10-week-old C3H/HeJ mice. Analysis of GSE45513 with the webtool GEO2R revealed expression of all 16 IGN genes in the mouse skin samples (Fig. 1).

**Fig. 1:**
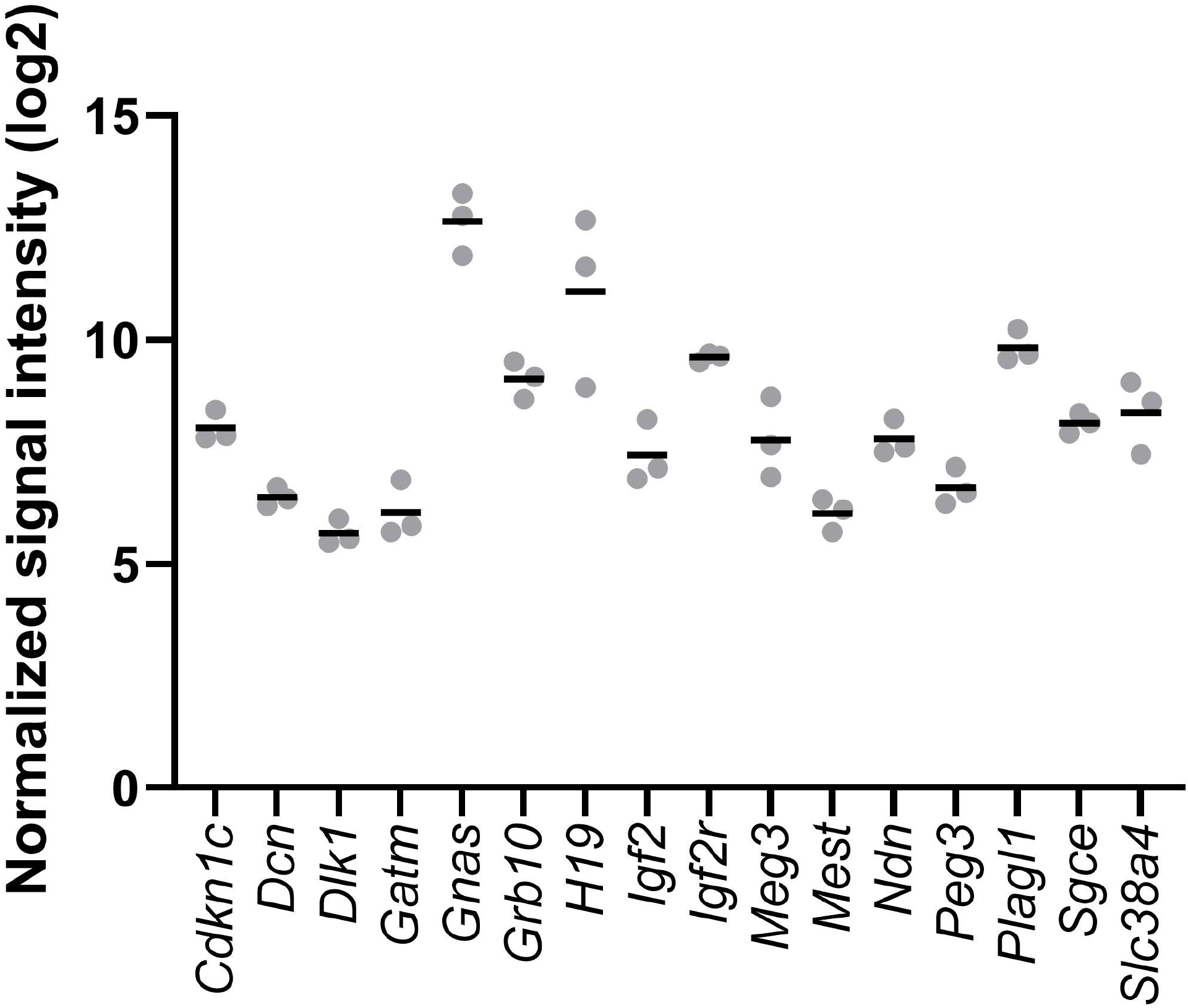
All 16 IGN genes are expressed in the dorsal skin of mice. Normalized signal intensity values (symbols) of all 16 IGN genes obtained from microarray data processed from skin samples of 10-week-old untreated and unaffected C3H/HeJ mice. The mean (horizontal line) of the normalized signal intensity values for each of the 16 IGN genes was calculated using the three biological repeats provided in GSE45513.

To identify publicly available datasets in which members of the IGN could be examined in the skin, we searched for one IGN gene *H19* using the search term ‘H19[gene symbol], AND skin’ in the NCBI GEO Profiles database ^17^. In this search, 156 datasets were identified. These datasets were manually curated for differential expression of *H19* across all samples within a dataset based on the visual gene expression level displayed in the GEO Profiles and using the GEO2R tool. Using this strategy, we identified dataset GSE11186 ^18^, which contains the transcriptomic profiles of the different stages of the first and second synchronized natural and depilation-induced growth cycles of hair follicles from mouse skin biopsies analyzed by Affymetrix array hybridization. The time-points representing the different phases of the synchronized hair growth cycle were classified by Lin *et al*. based on established morphological guidelines ^19^. In this dataset, *H19* expression was significantly elevated in the second telogen phase (day 44) compared to the mid-anagen (day 27) or catagen (days 37 and 39) phases (Fig. 2). As dataset GSE11186 contained only two samples for the first telogen phase (day 23), it was not included in this analysis

**Fig. 2.**
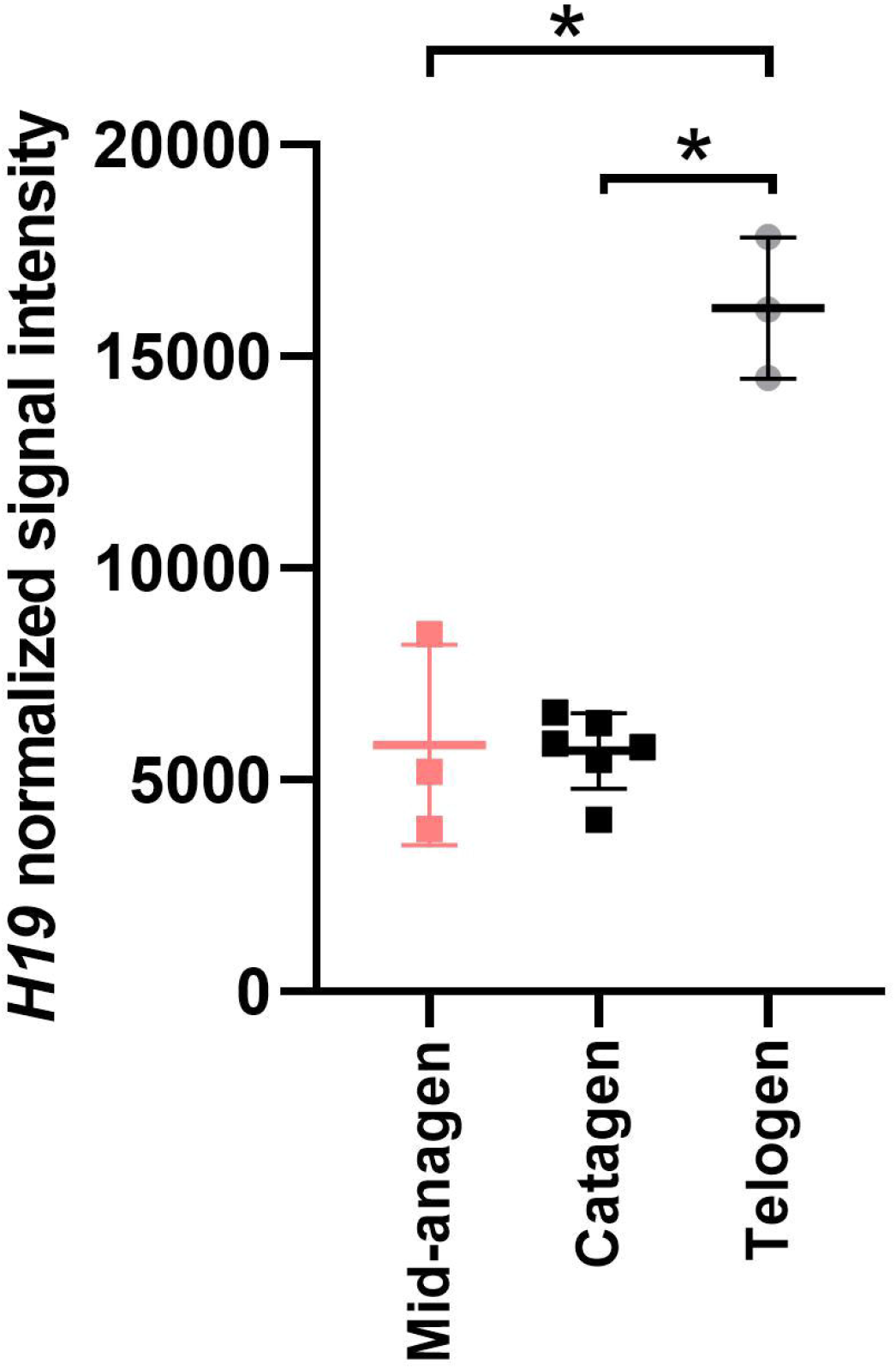
Time-course profile of *H19* expression during the synchronized second postnatal hair growth cycle. Values shown are quantile normalized signal intensity data from GSE11186 for the mid-anagen (day 27, n = 3), catagen (days 37 and 39, n = 6) and telogen (day 44, n = 3) phases. *P*-values shown were determined by Kruskal–Wallis analysis and Dunn’s test was used to determine which groups had significant differences *: *P* ≤ 0.05; n: number of samples provided in GSE11186 and used in the analysis.

This result prompted us to analyze the gene expression of all 16 IGN genes in the complex architecture of the hair follicle. Using the query ‘skin hair follicle’ to search the GEODataSet database, we identified dataset GSE3142, which contains the expression profiles of dermal papilla cells, skin fibroblasts, melanocytes, hair follicle matrix cells and outer root sheath cells from the dorsal skin of 4-day-old CD-1 mice ^20^, which represents the initial hair follicle morphogenesis stage. Our analysis revealed expression of all 16 IGN genes in the studied cell fractions of the hair follicle (Supplemental Fig. S1).

### 3.2 IGN gene expression is elevated during the telogen phase of the hair cycle

*H19* belongs to a network of 16 imprinted genes (*Cdkn1c, Dcn, Dlk1, Gatm, Gnas, Grb10*, H19, *Igf2, Igf2r, Meg3, Mest, Ndn, Peg3, Plagl1, Sgce*, and *Slc38a4*) co-expressed as part of an IGN that is regulated at the transition from proliferation to quiescence ^1,5^. *H19* knockout was shown to perturb the expression of five other IGN genes (*Igf2, Cdkn1c, Gnas, Dlk1* and *Igf2r*) at the transcriptional levels ^6^. Thus, in addition to *H19*, we next examined the normalized signal intensity values of each of the 16 individual IGN genes during the telogen (day 23) and mid-anagen (day 27) phases of the synchronized second postnatal hair cycle in the GSE11186 dataset. For comparison, we also examined the absolute expression of six known telogen-activated genes (*Ar, Esr1, Lhx2, Nr1d1, Sox18*, and *Stat3*), and six known telogen-repressed genes (*Elf5, Foxn1, Grhl1, Lef1, Msx2*, and *Vdr*) ^18^. We observed that the median IGN gene expression in the telogen phase was elevated compared to that in the mid-anagen phase. This trend was less marked for *Dcn* and *Igf2r*. Some of the IGN genes (i.e., *Gnas, H19, Meg3* and *Plagl1*) were expressed at even higher levels than the known telogen-activated genes (Fig. 3A).

**Fig. 3.**
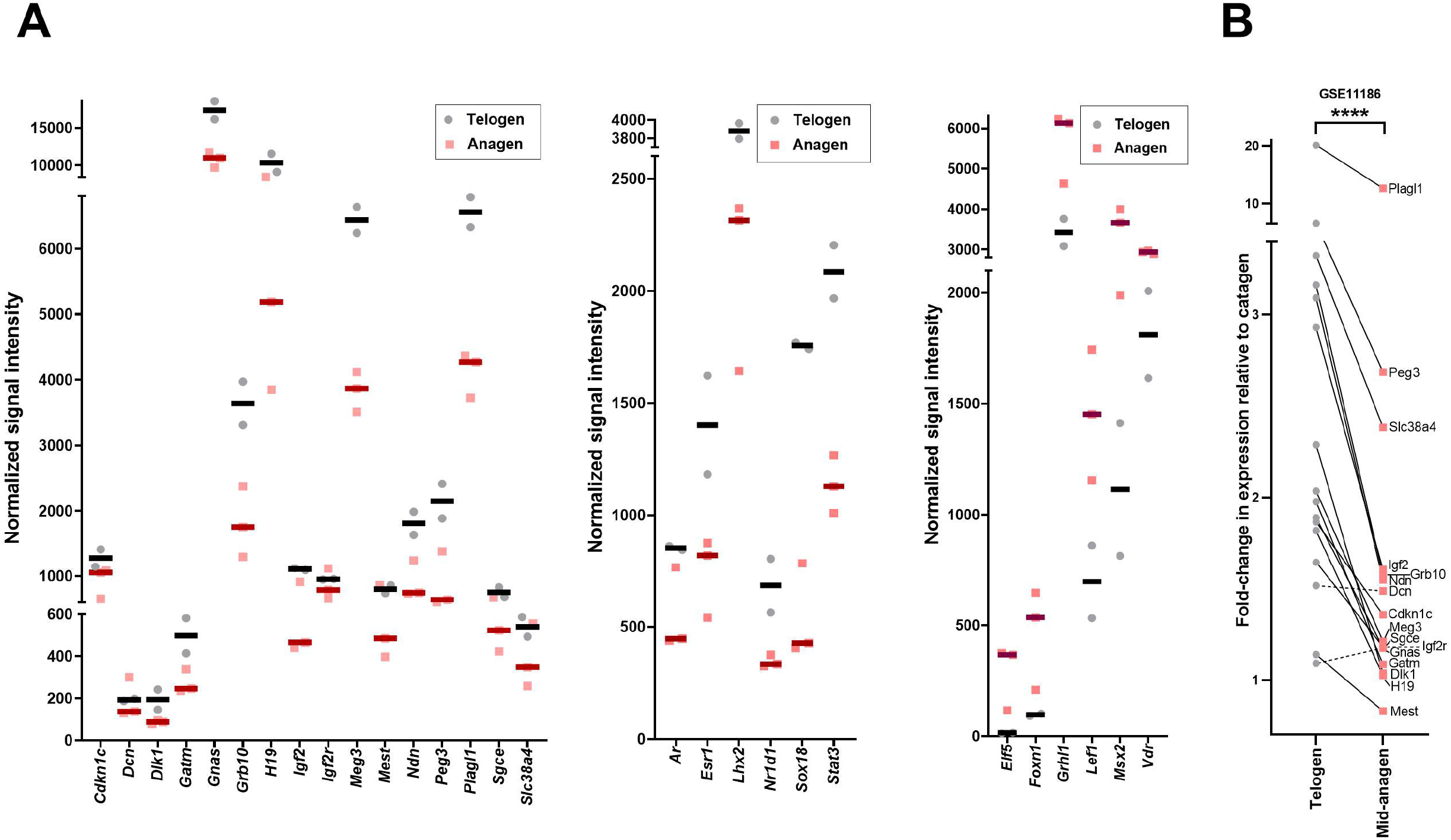
Expression of all 16 IGN genes in the telogen and anagen phases. (A) Dot plot showing the quantile normalized signal intensity data for the IGN and control genes in the telogen (day 23) and anagen (day 27) phases in GSE11186. Each dot (telogen: gray and anagen: red) corresponds to a subject from the GSE11186 dataset. IGN genes are more commonly expressed in the telogen phase compared to the anagen phase. Known telogen-activated genes (*Ar, Esr1, Lhx2, Nr1d1, Sox18*, and *Stat3*) and telogen-repressed genes (*Elf5, Foxn1, Grhl1, Lef1, Msx2*, and *Vdr*) ^18^ were used as controls. (B) Fold-change in expression of all 16 pooled IGN genes identified in the telogen phase (T, day 23) and the mid-anagen phase (A, day 27) normalized to that in the catagen phase (C, day 37 and day 39) in the GSE11186 dataset. T/C- and A/C-values that correspond to the same gene are connected with a line to demonstrate that the A/C fold-change in expression is lower than the T/C fold-change in expression for all IGN genes except *Dcn* and *Igf2r*. Statistical comparison of T/C- and A/C-values was performed using the Wilcoxon test, ****: *P* < 0.01.

Next, we calculated the fold-change in gene expression during the telogen phase (day 23) compared with that in the catagen phase (days 37 and 39) and during the anagen phase (day 27) compared with that in the catagen (days 37 and 39) phase using the normalized signal intensity values for each IGN member gene provided in GSE11186. We found a significant difference in the mean expression ratio of 14 IGN member genes (*Cdkn1c, Dlk1, Gatm, Gnas, Grb10, H19, Igf2, Meg3, Mest, Ndn, Peg3, Plagl1, Sgce*, and *Slc38a4)* when comparing the telogen/catagen ratio versus the mid-anagen/catagen ratio (Fig. 3B). Two of the panel of 16 IGN genes (*Dcn* and *Igf2r*) did not follow this trend (Fig. 3B).

In summary, our analysis shows that like *H19*, IGN genes are, in general, expressed at higher levels in the telogen phase compared to those in the anagen phase.

### 3.3 Most IGN genes are expressed periodically and are considered hair cycle-regulated genes

To assess whether the IGN genes are hair cycle-regulated, we took advantage of a publicly available dataset that was obtained after processing mouse skin mRNA microarray data obtained at eight time-points corresponding to the first synchronous (days 1, 6 and 14: anagen phase, day 17: catagen, day 23: telogen) and asynchronous (9^th^ week, 5^th^ month, 1^st^ year) periods of postnatal hair cycling ^21^. While the skin patches of the synchronous samples were collected at defined hair growth stages (anagen, catagen, telogen), the samples obtained during the asynchronous periods contain skin tissue at different phases of the hair cycle ^21^. Skin samples from synchronized and asynchronized hair cycle stages were included in order to distinguish changes in gene expression associated specifically with the hair cycle from non-cyclic changes in expression occurring simultaneously in the skin ^21^. After excluding genes that were not expressed in the mouse skin and applying a computational approach including replicate variance analysis (*F*-test), Lin *et al*. identified a dataset of 2,461 probe sets corresponding to 2,289 potential hair cycle-associated genes (hereafter referred to as the Lin1-dataset; Table S1) ^21^. The *P*-value cut-off for the *F*-test was set previously to 0.05, as it was found that >80% of known genes exhibiting hair cycle-dependent expression had a *P*-value of <0.05 determined using this computational approach ^21^. As the pool of these 2,289 hair cycle-associated genes was restricted to protein-coding genes, only the 14 protein-coding IGN genes (*Igf2, Cdkn1c, Dcn, Dlk1, Gatm, Gnas, Grb10, Igf2r, Ndn, Mest, Peg3, Plagl1, Sgce* and *Slc38a4*) were included in our analysis and the two non-coding RNA IGN genes (*H19* and *Meg3*) were excluded. Among the pool of 2,461 probe sets categorized as periodically expressed, hair cycle-regulated genes in mouse dorsal skin, we identified 10 IGN genes (*Cdkn1c, Dcn, Dlk1, Gatm, Gnas, Igf2r, Ndn, Peg3, Sgce*, and *Slc38a4*), corresponding to 71% of all protein-coding IGN genes (Fig. 4A). The previous cluster analysis of the Lin1-dataset revealed three general expression profile patterns, characterized as ‘anti-hair growth’-, ‘hair growth’-, and ‘catagen’-related, which were further subdivided into 30 sub-clusters of co-expressed genes with expression peaks at different stages of the hair cycle ^21^. Seven of the hair cycle-associated IGN genes (*Cdkn1c, Dlk1, Gnas, Peg3, Gatm, Ndn*, and *Slc38a4*) identified in our study were grouped in the ‘anti-hair growth’ category and showed a decline in expression levels during the anagen phase. *Igf2r* was categorized as a ‘hair growth’ gene, with peak expression early in the anagen phase. In contrast, *Sgce* was categorized as a ‘catagen-related’ gene, with a decrease in expression during the catagen phase. Only one IGN gene (*Dcn*) belonged to a gene cluster that could not be categorized according to the three main profile patterns (Table 1).

**Fig. 4.**
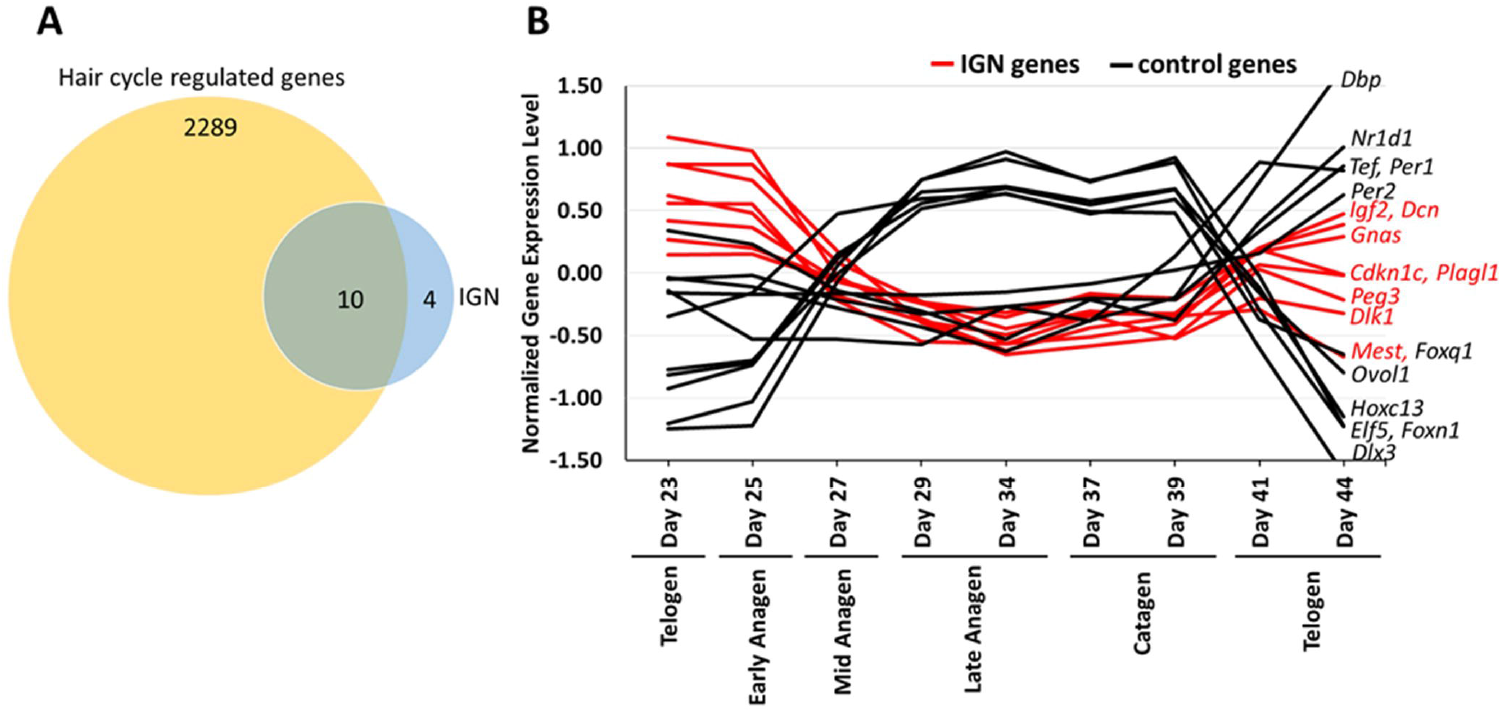
Most IGN genes are periodically expressed, hair growth cycle-regulated genes. **(A)** Venn diagram illustrating that 71% of protein-coding IGN genes (10 of 14) are hair cycle-regulated. The 10 protein-coding, hair cycle-regulated IGN genes identified are *Cdkn1c, Dcn, Dlk1, Gatm, Gnas, Igf2r, Ndn, Peg3, Sgce*, and *Slc38a4*. **(B)** Time-course profiles of hair cycle-regulated IGN gene expression during hair growth. The normalized expression levels of eight hair cycle-regulated IGN genes [(*Igf2, Cdkn1c, Dcn, Dlk1, Gnas, Mest, Peg3*, and *Plagl1*) are shown in red, and 11 control genes (telogen upregulated genes ^18^: *Dbp, Tef, Nr1d1, Per1, Per2*, and anagen/catagen upregulated genes ^18^: *Dlx3, Elf5, Foxn1, Foxq1, Hoxc13*, and *Ovol1*) are shown in black]. The y-axis represents the log-transformed, zero-mean gene expression (normalized gene expression level). Log-transformed, zero-mean gene expression values are provided in supplemental Table S1 ^18^.

**Table 1.** Potential hair cycle-associated IGN genes (identified in Lin1-dataset)

| Gene symbol | Gene name | P-value* | Cluster description* |
| --- | --- | --- | --- |
| *Cdkn1c* ** | Cyclin-dependent kinase inhibitor 1C (P57) | 0.0004 | Anti-hair growth pattern with decline in expression level during anagen phase |
| *Dcn* ** | Decorin | 0.0392 | Does not correspond with any of the three main profile patterns |
| *Dlk1* ** | Delta-like 1 homolog (*Drosophila*) | 0.0166 | Anti-hair growth pattern with decline in expression level during anagen phase |
| *Gatm* | Glycine amidinotransferase (L-arginine:glycine amidinotransferase) | 0.0009 | Anti-hair growth pattern with decline in expression level during anagen phase |
| *Gnas* ** | GNAS (guanine nucleotide binding protein, alpha stimulating) complex locus | 0.0040 | Anti-hair growth pattern with decline in expression level during anagen phase |
| *Igf2R* | Insulin-like growth factor 2 receptor | 0.0004 | Hair growth pattern including genes with expression peak early during anagen phase |
| *Ndn* | Necdin | 0.0171 | Anti-hair growth pattern with decline in expression level during anagen phase |
| *Peg3* ** | Paternally expressed gene 3 | 0.0001 | Anti-hair growth pattern with decline in expression level during anagen phase |
| *Sgce* | Sarcoglycan, epsilon | 0.0117 | Catagen-related expression patterns with decline in expression during catagen phase |
| *Slc38a4* | Solute carrier family 38, member 4 | 0.0013 | Anti-hair growth pattern with decline in expression during anagen phase |

\* Defined by Lin *et al.* <sup>21</sup>, \*\* Gene was independently identified as a potential hair cycle-regulated gene in the Lin2-dataset (see Table 2)

Next, we analyzed an independent dataset (previously reported by Lin *et al*. and hereafter referred to as the Lin2-dataset; Table S2), which comprises a set of 6,393 mRNA probe sets and corresponds to a pool of 4,704 genes ^18^ identified by processing expression data obtained from mRNA profiles of mouse dorsal skin collected at multiple time-points during: 1) the postnatal completion of hair follicle morphogenesis, including the first catagen and telogen phases; 2) the synchronized second postnatal hair growth cycle; and 3) a depilation-induced hair growth cycle ^18^. By applying a matrix model, 8,433 periodically expressed probe sets (6,010 genes) were identified, of which 2,040 (1,306 genes) were excluded from this subset since the changes in the expression of these genes was due to alterations in the cell type composition of the skin during hair growth (such as cornified cells, suprabasal cells, mesenchymal cells and myocytes) ^18^. The final set of 6,393 probe sets (4,704 genes, Lin2-dataset) exhibited periodic expression patterns that cannot be explained by cell type specific alterations that occur in the skin during hair growth and were thus defined as hair cycle-regulated genes ^18^. Similar to the Lin1-dataset, this set of 4,704 hair cycle-regulated genes was restricted to protein-coding genes. Thus, we included only the 14 protein-coding IGN genes in our analysis and excluded the two non-coding RNA IGN genes (*H19* and *Meg3*). We identified eight IGN genes (*Igf2, Cdkn1c, Dcn, Dlk1, Gnas*, Mest, *Peg3*, and *Plagl1*) (corresponding to 57% of all the protein-coding IGN genes) among the pool of 6,347 probe sets in the Lin2-dataset that were categorized as periodically expressed, hair cycle-regulated genes in mouse dorsal skin (Table 2) ^18^. A total of 3,180 genes from the Lin2-dataset were grouped previously according to their expression peak during the hair growth cycle, with 1,169 genes in the early anagen phase, 1,017 in the mid-anagen phase, 243 in the late anagen phase, 208 in the early catagen phase, 253 in the mid-catagen phase and 290 in the telogen phase ^18^. The eight IGN genes identified in this study that were included in the hair cycle-regulated genes of the Lin2-dataset were categorized as genes with an expression peak in the telogen phase (*Dcn*, and *Gnas*) and the early anagen phase (*Igf2, Cdkn1c, Dlk1, Mest, Peg3*, and *Plagl1*) (Table 2). Furthermore, we examined the expression levels of the eight hair cycle-regulated IGN genes (listed in Table 2) during the nine time-points provided in the ‘Lin2-dataset’^18^. The eight hair cycle-regulated IGN genes show elevated expression profiles in the telogen and early anagen phases compared to the mid/late anagen and catagen phases (Fig. 4B). This expression pattern is similar to that of *Dbp, Nr1d1, Per1, Per2*, and *Tef*, which form a co-expressed cluster of transcription factors known to be elevated during the telogen phase ^18^. In contrast, their expression patterns differed from those of *Dlx3, Elf5, Foxn1, Foxq1, Hoxc13*, and *Ovol1*, a group of key transcriptional regulators with known peaks in expression from the mid-anagen to late catagen phase ^18^. In summary, our analysis shows that the majority of IGN genes are among two independently identified datasets ^18,21^ containing hair cycle-regulated genes, the expression of which is not linked to cell type specific alterations that occur in the skin during the hair cycle. In addition, we identified the IGN genes among ‘Telogen’ and ‘Early Anagen’ genes grouped by Lin *et al*. using statistical differential analysis ^18^.

**Table 2.**
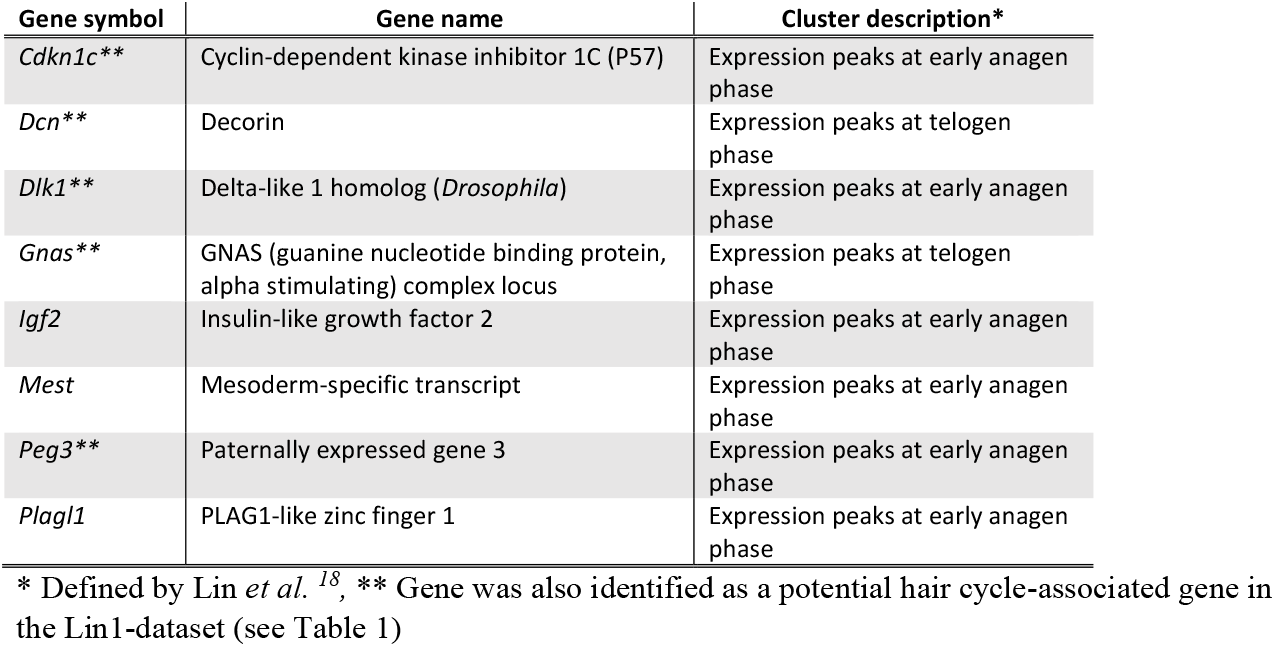
Potential hair cycle-regulated IGN genes (identified in Lin2-dataset)

### 3.4 Network analysis reveals a potential role of IGN genes as upstream regulators of hair cycle-associated genes

Hair follicle development and regeneration *in vitro* are strictly regulated by various growth factors, hormones and signaling molecules, with the Wnt signaling pathway being one of the most important ^22-26^. A study reported by Zhu *et al*. indicated that H19 maintains the hair follicle-inducing ability of dermal papilla cells through activation of the Wnt pathway ^27^.Therefore, we explored the potential function of the 16 IGN genes as upstream regulators of hair cycle-associated genes. For this purpose, we performed IPA using the hair cycle-associated genes listed in Table S1 as input to identify potential upstream regulators of these genes. We successfully identified eight IGN genes (*Cdkn1c, Dcn, Gnas, Grb10, H19, Igf2, Igf2r*, and *Plagl1*) among the predicted upstream regulators of hair cycle-associated genes (Supplemental Fig. S2, Table S3).

To further explore the relationships of all 16 IGN genes and to identify potentially associated biological functions, we used the IPA application to perform a network analysis with all 16 IGN genes as input. Using this strategy, we identified a larger gene network comprising 35 genes, of which 12 were IGN genes (*Cdkn1c, Dcn, Dlk, Gatm, Gnas, Grb10, H19, Igf2, Igf2r, Meg3, Ndn, Plagl1*) (Table S4, Fig. 5). In the IPA, this network of genes was predicted to be associated with ‘Dermatological Diseases and Conditions’ in the ‘Top Diseases and Functions’ category. To explore potential functions of this network further, we sought to determine whether these 35 genes were among the hair cycle-associated genes listed in Table S1. Our analysis revealed that 10 genes from this network had previously been identified as hair cycle-associated genes (Table S4). Further analysis revealed that 27 of the 35 genes in the larger IGN network were among the potential upstream regulators of the hair cycle-associated genes listed in Table S3 (Table S4 and Fig. 5).

**Fig. 5.**
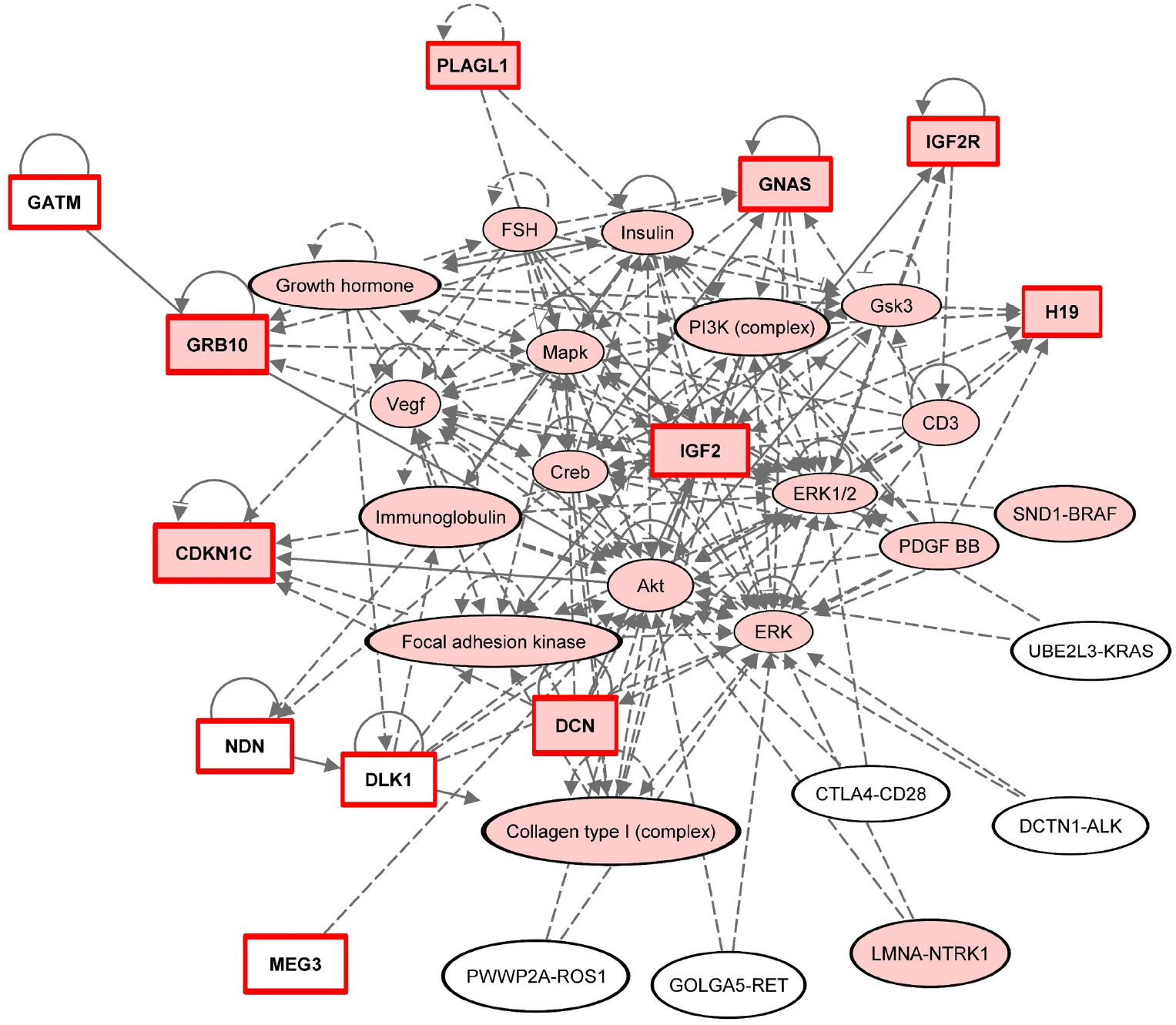
IPA reveals a network including IGN genes. IPA revealing the annotated interactions of 35 genes including 12 IGN genes (*Cdkn1c, Dcn, Dlk1, Gatm, Gnas, Grb10, H19, Igf2, Igf2r, Meg3, Ndn*, and *Plagl1*, indicated by a square box with a red outline and bold text). The symbols for the 26 genes identified as potential upstream regulators of hair cycle-associated genes are filled in light red.

Finally, we explored potential downstream functions of the 16 telogen/early anagen-activated IGN genes using the canonical pathway analysis tool in the IPA application. We identified 10 canonical pathways in which seven (*Dcn, Gatm, Gnas, H19, Igf2, Igf2r*, and *Ndn*) of the 16 IGN genes were significantly enriched based on a -log (*P*-value) >1.3 (Table 3). These canonical pathways included hormone signaling pathways, such as the estrogen receptor pathway, which was previously shown to regulate the transition from the telogen phase to the anagen phase in hair follicles ^28^, as well as several metabolism-and signaling-related pathways such as the glycine degradation (creatine biosynthesis) pathway and ephrin B signaling (Table 3).

**Table 3.** Identified canonical pathways with the 16 IGN genes significantly enriched.

| Ingenuity canonical pathways | -log(P-value) | Ratio | z-score | IGN-enriched genes |
| --- | --- | --- | --- | --- |
| Glycine degradation (creatine biosynthesis) | 2.85 | 0.5 | 0 | *Gatm* |
| Estrogen receptor signaling | 2.84 | 0.00915 | 0 | *Gnas, Igf2, Igf2r* |
| Glioma signaling | 2.58 | 0.0182 | 0 | *Igf2, Igf2r* |
| White adipose tissue browning pathway | 2.44 | 0.0155 | 0 | *Gnas, Ndn* |
| Osteoarthritis pathway | 2.03 | 0.00948 | 0 | *Dcn, H19* |
| Apelin muscle signaling pathway | 1.88 | 0.0526 | 0 | *Gnas* |
| G protein signaling mediated by Tubby | 1.67 | 0.0323 | No activity pattern available | *Gnas* |
| PFKFB4 signaling pathway | 1.5 | 0.0217 | 0 | *Gnas* |
| Growth hormone signaling | 1.31 | 0.0141 | 0 | *Igf2* |
| Ephrin B signaling | 1.31 | 0.0139 | 0 | *Gnas* |

## 4. Discussion

The 16 IGN genes examined in this study are known to play important roles in embryonic development ^5,29^. In this study, we investigated the involvement of this particular set of imprinted genes in the growth cycle of hair follicles and the underlying mechanisms that regulate the expression of hair cycle-associated genes. We identified a coordinated elevation in the expression of these IGN genes in the telogen and early anagen phases compared to the mid-anagen phase of the hair cycle. In addition, we identified most IGN genes among a list of previously reported hair cycle-associated genes. Our analysis showed that IGN genes form a network with other genes, most of which most were characterized as upstream regulators of hair cycle-associated genes. In addition, our gene enrichment analysis using IPA predicted eight of the 16 IGN genes as potential upstream regulators of hair cycle-associated genes. Thus, our findings indicate a potential novel role of IGN genes as upstream regulators of hair cycle-associated genes.

Patterns of gene expression in hair follicle stem cells have been reported previously ^30,31^ as well as time-course profiling of the expression in the skin to identify hair cycle-associated genes, including the dataset reported by Lin *et al*. that was used for the analysis in our study ^21^. However, synchronous, coordinated hair cycle-associated changes in the expression of IGN genes have not been described previously. Our analysis of the cyclic expression of IGN genes in hair follicles as well as their potential functions as upstream regulators of hair cycle-regulated genes highlight the vital role of IGN genes in skin/hair homeostasis ^7^.. Interestingly, a recent study has shown that *H19* overexpression activates the Wnt signaling pathway, resulting in maintenance of the hair follicle regeneration potential ^32^. This activity of H19 may, to some extent, contribute to the molecular mechanisms underlying our findings indicating that *H19* and other IGN genes function as potential upstream regulators of hair cycle-regulated genes.. In addition, Iglesias–Bartolome *et al*. showed that the *GNAS* gene product, which we identified as a potential telogen-activated upstream regulator of hair cycle-regulated genes, limits the proliferation of epidermal stem cells and is involved in maintaining hair follicle homeostasis ^33^. Moreover, we identified *Dcn*, a member of the IGN, as an upstream regulator of hair cycle-regulated genes (see Table S3 and Fig. 5). Exogenous administration of its gene product, decorin, was shown to accelerate the anagen phase and delay catagen phase transition, and was categorized as a positive regulator of the hair growth cycle ^34^. It has also been shown that physiologic concentrations of *Igf2*, which is among our predicted upstream regulators of hair cycle-associated genes (Table S3), is a potent stimulator of hair growth ^35^. Furthermore, absence of *Igf2* results in premature entry into a catagen-like stage ^35^. Thus, in accordance with our conclusions, *Igf2* was suggested to be an important regulator of the hair cycle ^35^.

Interestingly, most genes in the larger network of regulatory IGN genes identified in this study (27 of 35 genes) were identified as potential upstream regulators of hair cycle-associated genes (Table S3, S4 and highlighted in red in Fig. 5). Among the genes predicted to be directly regulated by the IGN, we identified several known to be associated with regulation of the hair cycle, including *Akt*, Erk, *Pi3k*, and *Vegf* (Fig. 5). Indeed, the PI3K-Akt signaling pathway plays a crucial role in *de novo* hair follicle regeneration ^36^, while vascular endothelial growth factor (*Vegf*) induces proliferation of hair follicle cells through activation of ERK ^37^.

The IGN genes examined in this study are downregulated postnatally, but are constitutively expressed in pluripotent stem cells and/or progenitor cells of the hematopoietic system, skin and skeletal muscles, with significantly lower expression levels in their differentiated progeny ^7^. Somatic stem cells, such as hematopoietic stem cells, epidermal stem cells and satellite cells of the skeletal muscles are generally considered to be quiescent, dividing infrequently, but are driven into active proliferation/differentiation cycles during tissue regeneration or self-renewal ^7^. Our results suggest periodic expression of the IGN genes during the follicular growth cycle, with peak expression in the telogen and early anagen phases and the lowest expression in the mid-anagen and catagen phases. These findings are consistent with the consensus that hair follicle stem cells receive activating or inhibitory signals at distinct stages of the hair growth cycle, allowing them to either remain quiescent or become proliferative ^38^. During the transition from the telogen phase to the anagen phase, biological signals from the dermal papilla stimulate the quiescent follicular stem cells to proliferate ^39^. Melanocytes, which are important components of hair follicles that produce hair color, are also activated at specific phases of the hair growth cycle, supplying progeny to the hair matrix, where most mature into differentiated melanocytes ^40,41^. As most of the IGN genes have the potential to function as tumor suppressors, it is likely that the decrease in their expression from the telogen phase to the anagen phase triggers biological cascades that stimulate cell proliferation in melanocytes ^42-52^. Interestingly, H19 downregulation was shown to stimulate melanogenesis in melasma, a hyperpigmentation condition resulting from an increase in melanin pigment production ^53^. Moreover, it can be speculated that dysregulated IGN gene expression plays roles in the development of some of the manifestations of several congenital syndromes characterized by impaired hair growth cycles, including short anagen hair syndrome ^54^, which is associated with a synchronized pattern of scalp hair growth, ^55^ and androgenic alopecia, which is a very common type of hair-loss ^32^. As H19 overexpression was shown to activate Wnt signaling to maintain the hair follicle regeneration potential, it has already been suggested that H19 could be a target for treatment of androgenetic alopecia ^27^.

The limitations of our study should be noted. Our study was designed to examine the potential function of the 16 IGN genes in mouse skin tissue after birth by reanalyzing publicly available transcription profiles. Thus, this study consists of a *in silico* analysis of these 16 IGN genes, but does not include wet-lab characterization and functional analysis. Not only is the hair follicle a complex mini-organ that presents some challenges to wet-lab investigations, but the complementary and contrasting functions of the proteins encoded by the IGN genes represent a challenge in determining their individual functions in the hair cycle. In-depth studies might require the simultaneous knockout of different combinations of the coordinately expressed network genes as a single gene might not alter the integrity of the entire network. Nevertheless, we consider that our *in silico* analysis of several independent datasets supports the conclusion that IGN genes are periodically expressed in a coordinated manner during the hair cycle and might participate in syndromes characterized by an impaired hair cycle when dysregulated.

In summary, we have shown that the majority of genes belonging to the IGN show synchronous, coordinated expression in mouse skin during the hair cycle, with elevated expression in the telogen and early anagen phases. In addition, we revealed that IGN genes form part of a larger network including non-imprinted genes together may function as upstream regulators of hair cycle-regulated genes. Based on our findings, we propose a novel role for IGN genes in regulating progression of the hair cycle. Our observation that IGN genes are more abundantly expressed in the telogen and early anagen phases indicates a possible role for IGN genes in the control of this stage of the hair cycle.

## 5. Materials and Methods

### Dataset search and analysis

To identify publicly available transcription profile data of untreated and unaffected mouse skin, we used the query ‘skin AND C3H/HeJ’ to search the NCBI GEO DataSets, a public genomics data repository ^56^. We included the search term C3H/HeJ, as this is a general-purpose strain of mice used in a wide variety of research areas. Using this strategy, we identified GSE45513, which contains skin expression profiles of three 10-week-old untreated and unaffected C3H/HeJ mice ^57^. This dataset is provided as normalized signal intensity (log2) values. For the analysis of IGN gene expression in GSE45513, we used the interactive webtool GEO2R ^58^; see ‘Dataset analyses using GEO2R’. The following probes were used to evaluate expression of the genes indicated: 1417649_at (*Cdkn1c*), 1441506_at (*Dcn*), 1449939_at (*Dlk1*), 1423569_at (*Gatm*), 1450186_s_at (*Gnas*), 1425458_a_at (*Grb10*), 1448194_a_at (*H19*), 1448152_at (*Igf2*), 1424112_at (*Igf2r*), 1452905_at (*Meg3*), 1423294_at (*Mest*), 1415923_at (*Ndn*), 1417356_at (*Peg3*), 1426208_x_at (*Plagl1*), 1420688_a_at (*Sgce*), and 1428111_at (*Slc38a4*). The mean of the normalized signal intensity values of all three biological repeats for each IGN gene was calculated using GraphPad Prism (GraphPad Prism version 9.2.0 for Windows, GraphPad software, San Diego, CA USA).

To identify datasets in which our genes of interest (16 IGN genes) are differentially expressed, we used the query ‘H19[gene symbol] AND skin’ to search the NCBI GEO Profiles ^59^. This database stores gene expression data derived from the curated GEO datasets ^56^. Using our specific query, we identified expression profiles (presented as charts of transcriptomic datasets) containing samples with differentially expressed *H19* levels across all samples within each dataset. We then curated the filtered datasets based on differential gene expression of *H19* visually as well as using the NCBI GEO2R tool ^60^; see ‘Dataset analysis using GEO2R’. Using this strategy, we identified GSE1912 (Lin1 dataset) ^21^ and GSE11186 (Lin2 dataset) ^18^ which contain expression profiles of mouse dorsal skin at different stages of the synchronized and unsynchronized hair cycle. The expression profiles are provided as normalized signal intensity values (linear). The same probe IDs as listed above for the analysis of GSE45513 were used to analyze the Lin1 and Lin2 datasets.

To study the gene expression of the IGN genes in the different cell types of the hair follicle, we used the query ‘skin hair follicle’ in the GEODataSet database search, which was restricted to the organism ‘mouse’. This search revealed GSE3142 ^20^, a dataset that contains microarray expression profiles of hair follicle matrix cells, outer root sheath cells, dermal papilla cells, and melanocytes as well as a dermal fraction enriched in fibroblasts from the dorsal skin of 4-day-old CD-1 mice. Data values of GSE3142 are provided as normalized signal intensity values (log2). For the analysis of IGN gene expression in GSE3142, we downloaded the normalized signal intensity data (log2) and used GEO2R to determine the normalized signal intensity values for each of the 16 IGN genes in both biological replicates. The same probe IDs as listed above for the analysis of GSE45513 were used to analyze GSE3142. Finally, we calculated the mean of the two signal intensity values for each gene and created the superimposed scatter plot in GraphPad Prism 9.2.0.

Dataset analysis using GEO2R: After grouping the samples in GEO2R, the analysis was performed and dataset quality was assessed by the generated graphical plots provided in GEO2R. In brief, these comprised: 1) a volcano plot, generated using limma, displaying statistical significance versus magnitude of change to visualize differentially expressed genes; 2) a mean difference plot displaying log2 fold-change versus average log2 expression values to visualize differentially expressed genes; 3) a boxplot, generated using R boxplot, to view the distribution of the values of the selected samples; and 4) an expression density plot, generated using R limma, to view the distribution of the values in the selected samples. If dataset quality was satisfactory, the full table of normalized signal intensity values, including probe IDs and gene names, was downloaded. The probe IDs for each of the 16 IGN genes were determined. The probe IDs were entered sequentially into the search field in the Profile Graph tab in GEO2R and the normalized signal intensity values for the samples of interest were copied into GraphPad 9.2.0. to generate a graphical plot.

Statistical comparison of *H19* expression levels among three different growth phases of hair follicles was performed with the non-parametric Kruskal–Wallis test ^61^ and the Dunnett multiple comparison post-hoc test ^62^. The results were adjusted for multiple hypothesis testing with a Benjamini–Hochberg (fdr) procedure ^63^ Comparison of two ratios was performed with the non-parametric Wilcoxon test. Comparison of *Dcn* and *Gnas* expression between anagen and telogen from dataset GSE129218 was performed using the unpaired t-Test. A *P*-value ≤ 0.05 was considered statistically significant. Analyses were performed using R version 4.0.4.

### Skin tissue sample collection and microarray experiments

Dataset GSE4553 generation: The authors of GSE45513 isolated total RNA from skin samples of three 10-week-old untreated and unaffected C3H/HeJ mice. This was converted to cDNA and hybridized onto Mouse Genome 430 2.0 gene chips. Microarray quality control and preprocessing were performed by the authors of GSE45513 using Bio Conductor in R and RMA normalization.

Dataset GSE11186 generation: Dataset GSE11186 was generated by Lin *et al*. ^18^ in a microarray analysis of mouse dorsal skin samples obtained at postnatal time-points representative of the second synchronized hair cycle: day 23 (telogen), day 27 (mid-anagen), day 37–39 (catagen), and day 44 (telogen). Multiple biological replicates were profiled for each time-point (telogen day 23: n = 2; mid-anagen day 27: n = 3; catagen day 37–39: n = 6; telogen day 44: n = 3). In brief, fragmented biotinylated cRNA was hybridized on GeneChip Drosophila Genome Arrays. The authors of GSE11186 analyzed the microarray data with Microarray Suite version 5.0 (MAS 5.0) using Affymetrix default analysis settings and global scaling as the normalization method. Histological sections were used to classify each skin sample into specific phases/stages of the hair cycle based on established morphological guidelines ^19^.

Dataset GSE3142 was generated by Rendl *et al*. ^20^ in a microarray analysis of five cell types isolated from mouse dorsal skin samples. Briefly, Rendl *et al*. FACS-sorted hair follicle matrix cells, outer root sheath cells, dermal papilla cells, and melanocytes as well as a dermal fraction enriched in fibroblasts from dorsal skin samples from 4-day old Lef1-RFP/K14-H2BGFP mice. Two biological replicates of each cell type were generated. Total RNA isolated from each cell type was reverse-transcribed and biotinylated cRNA was hybridized to mouse genome array MOE 430a (Affymetrix). Microarray data processing was performed using the Affymetrix Microarray Suite version 5.0, scaled with the Affymetrix mask file set to TGT = 500.

Identification and classification of hair cycle-associated genes: Hair cycle genes were identified by Lin *et al*. ^21^ using data from a microarray analysis study including dorsal skin samples collected from CB6F1 mice at defined time-points during the synchronized stages of the hair cycle, as well as data from asynchronized skin samples. Histological sections were used to classify each sample of the synchronized stages of the hair cycle based on established morphological guidelines ^19^. Computational methods were then used to identify the set of genes expressed within the skin that are associated specifically with the hair growth cycle ^21^.

### Ingenuity pathway analysis

To identify potential upstream regulators, including transcription factors and any gene or small molecule that has been reported to affect gene expression experimentally, we used the web-based application Ingenuity Pathway Analysis (IPA, Ingenuity Systems Inc, Redwood City, CA, USA, version 62089861) ^64^. The upstream regulator analysis in the IPA application is based on prior knowledge of expected effects between transcriptional regulators and their target genes stored in the Ingenuity Knowledge Base ^65^. For this analysis, we uploaded the dataset of interest to the IPA application and performed a ‘Core Analysis’ using the Ingenuity Knowledge Base as a reference set and the ‘Upstream Regulator’ analytics. Similarly, the network analysis was generated using the 16 IGN genes as input data, performing a ‘Core analysis’ and using the ‘Networks’ analysis tool option in IPA. Using the same Core analysis of the 16 IGN genes, we also performed a canonical pathway analysis using the ‘Canonical Pathways’ tab in the IPA application. The resulting -log(*P*-values) were calculated by the IPA software using Fisher’s exact test to determine the probability that the association between the IGN genes and the identified canonical pathways is due to chance alone. A -log(*P*-value) of ≥1.3 was considered statistically significant.

## 7 Acknowledgments

The authors would like to thank Drs. D. Chaussabel, N. Marr, Dr. E. Chin-Smith and Jessica Tamanini for critically reading the manuscript. The authors would like to thank all the researchers who made their datasets public in the NCBI GEO.

## 8 Author contributions

AKM: conceptualization. SB, AKM: data curation, validation and visualization, data analysis and interpretation and methodology development. MT: data analysis. AKM: writing of the first draft. AKM, AIC, TK: writing, review and editing, MEA: statistical analysis. The contributor’s roles listed above follow the Contributor Roles Taxonomy (CRediT) managed by The Consortia Advancing Standards in Research Administration Information (CASRAI) (https://casrai.org/credit/). All authors contributed to the article and approved the submitted version.

## 11 Data availability

The datasets underlying the results are available in the NCBI GEO DataSets (GSE11186) at ncbi.nlm.nih.gov/gds/ as well as in the supplemental material of references ^18^ and of ^21^.

