## Supplementary figures and images for "In silico analysis of imprinted gene expression in the mouse skin"

### Figure S1

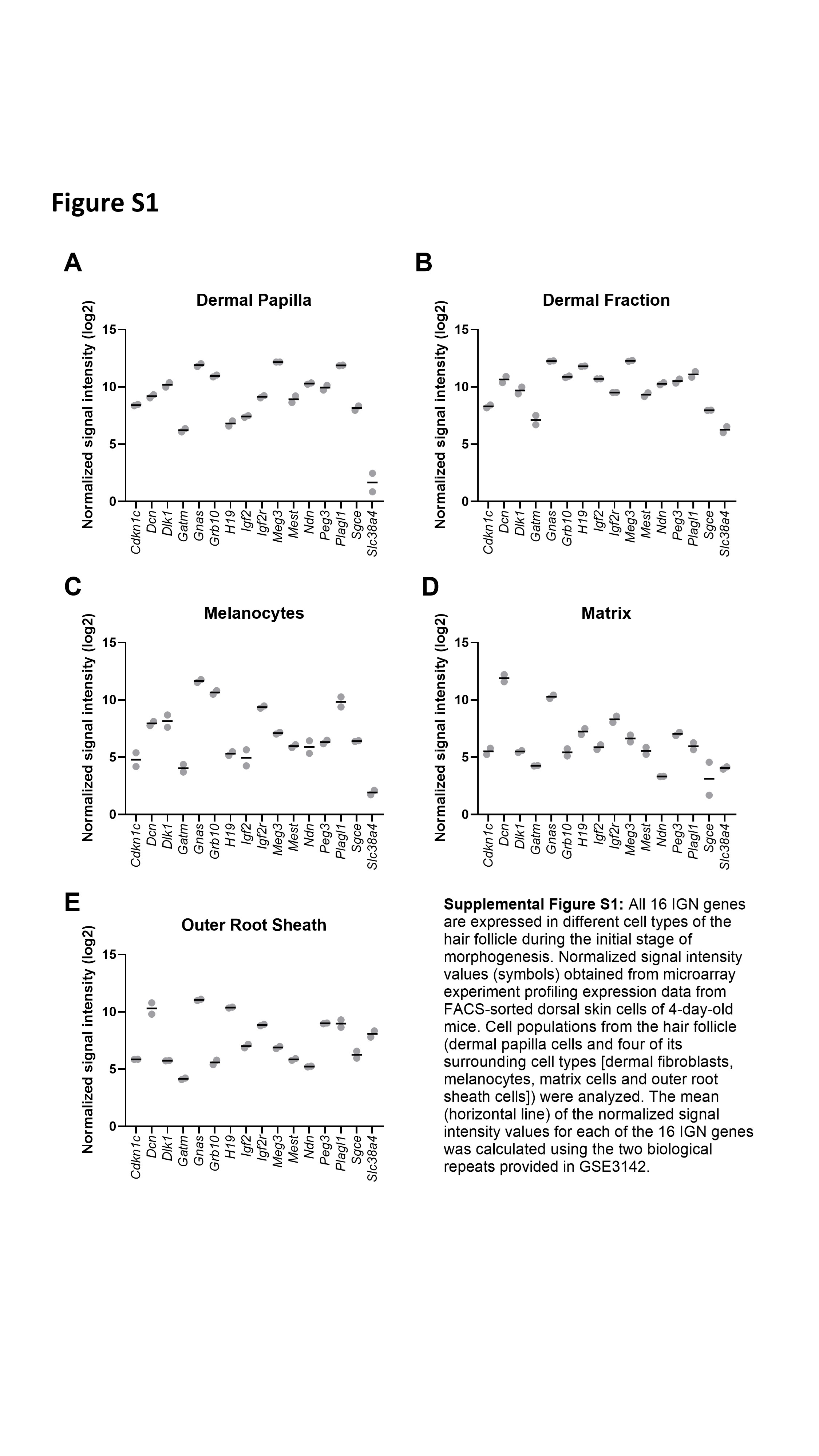

### Figure S2

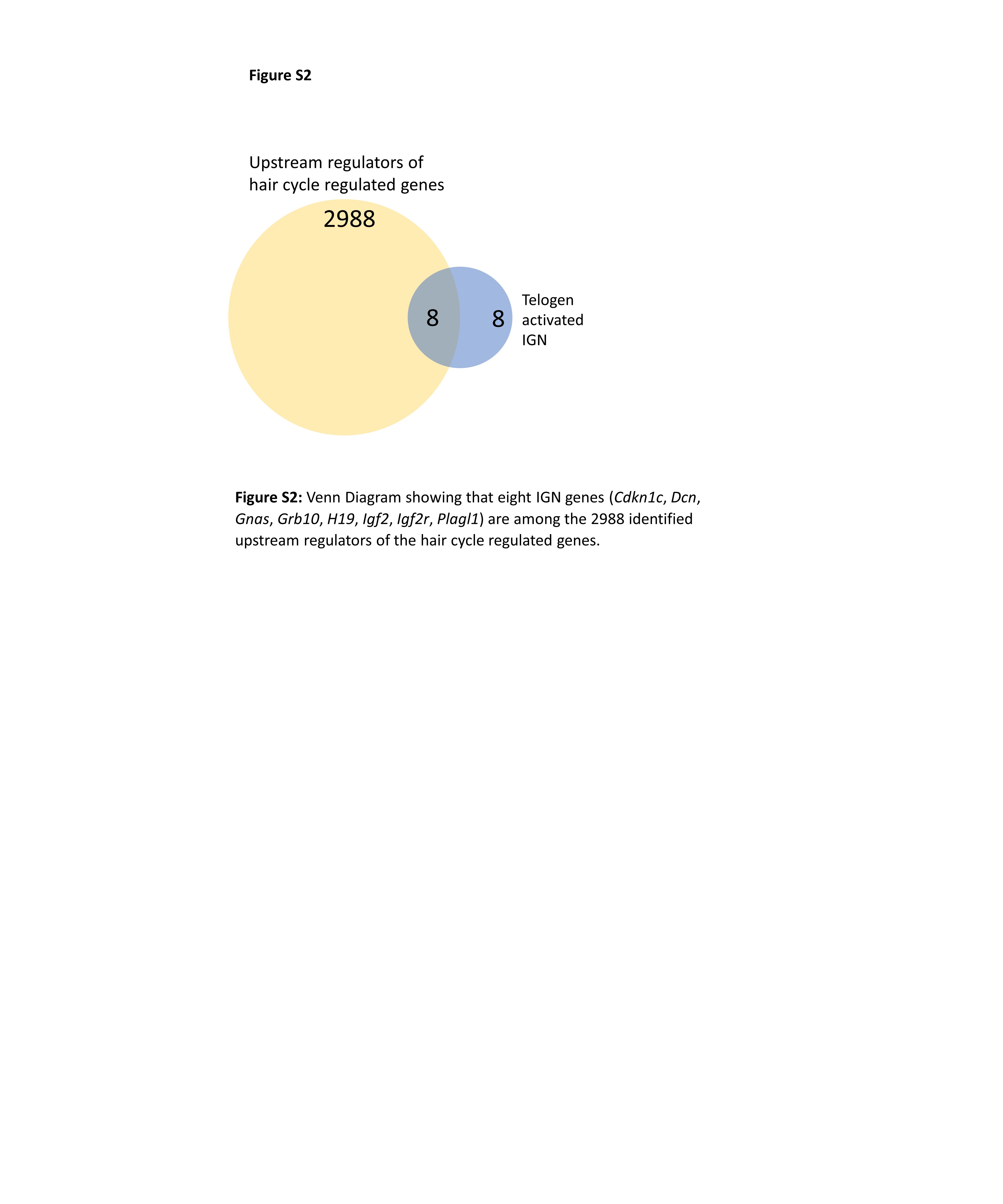
